# Biosynthesis of aurodox, a Type III secretion system inhibitor from *Streptomyces goldiniensis*

**DOI:** 10.1101/2022.02.14.480342

**Authors:** Rebecca E McHugh, John T Munnoch, Robyn E Braes, Iain J. W. McKean, Josephine M Giard, Andrea Taladriz-Sender, Frederik Peschke, Glenn A Burley, Andrew J Roe, Paul A Hoskisson

**Author notes:** **Supplementary Data:** McHugh et al., 2022 - Biosynthesis of aurodox, a Type III secretion system inhibitor from *Streptomyces goldiniensis*. Figshare. https://doi.org/10.6084/m9.figshare.19140005.v1.

## Abstract

The global increase in antimicrobial-resistant infections means that there is a need to develop new antimicrobial molecules and strategies to combat the issue. Aurodox is a linear polyketide natural product that is produced by *Streptomyces goldiniensis,* yet little is known about aurodox biosynthesis or the nature of the biosynthetic gene cluster (BGC) that encodes its production. To gain a deeper understanding of aurodox biosynthesis by *S. goldiniensis,* the whole genome of the organism was sequenced, revealing the presence of an 87 kb hybrid Polyketide Synthase/Non-Ribosomal Peptide Synthetase (PKS/NRPS) BGC. The aurodox BGC shares significant homology with the kirromycin BGC from *S. collinus* Tϋ 365; however, the genetic organisation of the BGC differs significantly. The candidate aurodox gene cluster was cloned and expressed in a heterologous host to demonstrate that it was responsible for aurodox biosynthesis and disruption of the primary PKS gene (*aurAI*) abolished aurodox production. These data support a model whereby the initial core biosynthetic reactions involved in aurodox biosynthesis follow that of kirromycin. Cloning *aurM\** from *S. goldiniensis* and expressing this in the kirromycin producer *S. collinus* Tϋ 365 enabled methylation of the pyridone group, suggesting this is the last step in biosynthesis. This methylation step is also sufficient to confer the unique Type III Secretion System inhibitory properties to aurodox.

**Importance:** Enterohaemorrhagic *Escherichia coli* (EHEC) is a significant global pathogen for which traditional antibiotic treatment is not recommended. Aurodox inhibits the ability of EHEC to establish infection in the host gut through the specific targeting of the Type III Secretion System, whilst circumventing the induction of toxin production associated with traditional antibiotics. These properties suggest aurodox could be a promising anti-virulence compound for EHEC, which merits further investigation. Here, we have characterised the aurodox biosynthetic gene cluster from *Streptomyces goldiniensis* and have established the key enzymatic steps of aurodox biosynthesis that give rise to the unique anti-virulence activity. These data provide the basis for future chemical and genetic approaches to produce aurodox derivatives with increased efficacy and the potential to engineer novel elfamycins.

## Introduction

*Streptomyces* bacteria are renowned for their ability to produce a plethora of natural products that exhibit a wide range of chemical structures, activities and modes of action (1). One such molecule is aurodox, which has a remarkable anti-virulence mode of action in addition to its well understood anti-gram-positive properties (2–4). Aurodox is produced by *Streptomyces goldiniensis* and belongs to the elfamycin group of antibiotics, which are characterised by their mode of action rather than their chemical structure (4). The anti-bacterial mode of action of the elfamycins is well understood, where they target protein translation through inhibition of Elongation Factor Thermo-unstable (EF-Tu; (4). Direct EF-Tu binding by kirromycin/aurodox-type elfamycins prevents EF-Tu:GDP from dissociating from the ribosome, preventing elongation and inhibiting protein synthesis (4). Aurodox also has an additional mode of action, originally discovered during a screen for Type III Secretion System (T3SS) inhibitors in Enteropathogenic *Escherichia coli* (EPEC; (5). More recently, it was demonstrated that aurodox inhibits T3SS and virulence in Enterohaemorrhagic *E. coli* (EHEC) and EPEC through an EF-Tu-independent mechanism, involving the downregulation of transcription of the master virulence regulator, Ler (6).

Aurodox was discovered in 1973 (2), it is a linear polyketide compound which is highly similar to kirromycin (7) differing only in methylation of the pyridone moiety. Kirromycin biosynthesis has been characterised and the BGC encodes five large polyketide synthase (PKS) units which act to form the polyketide backbone, before tailoring enzymes decorate the molecule (7–10). Given the similarity of the molecules, we hypothesised that the aurodox biosynthetic gene cluster (BGC) may be homologous to the hybrid PKS/Non-ribosomal peptide synthetase (NRPS) BGC of kirromycin, with the addition of an ORF responsible for pyridone-associated methylation (**Fig. 1**).

**Figure 1:**
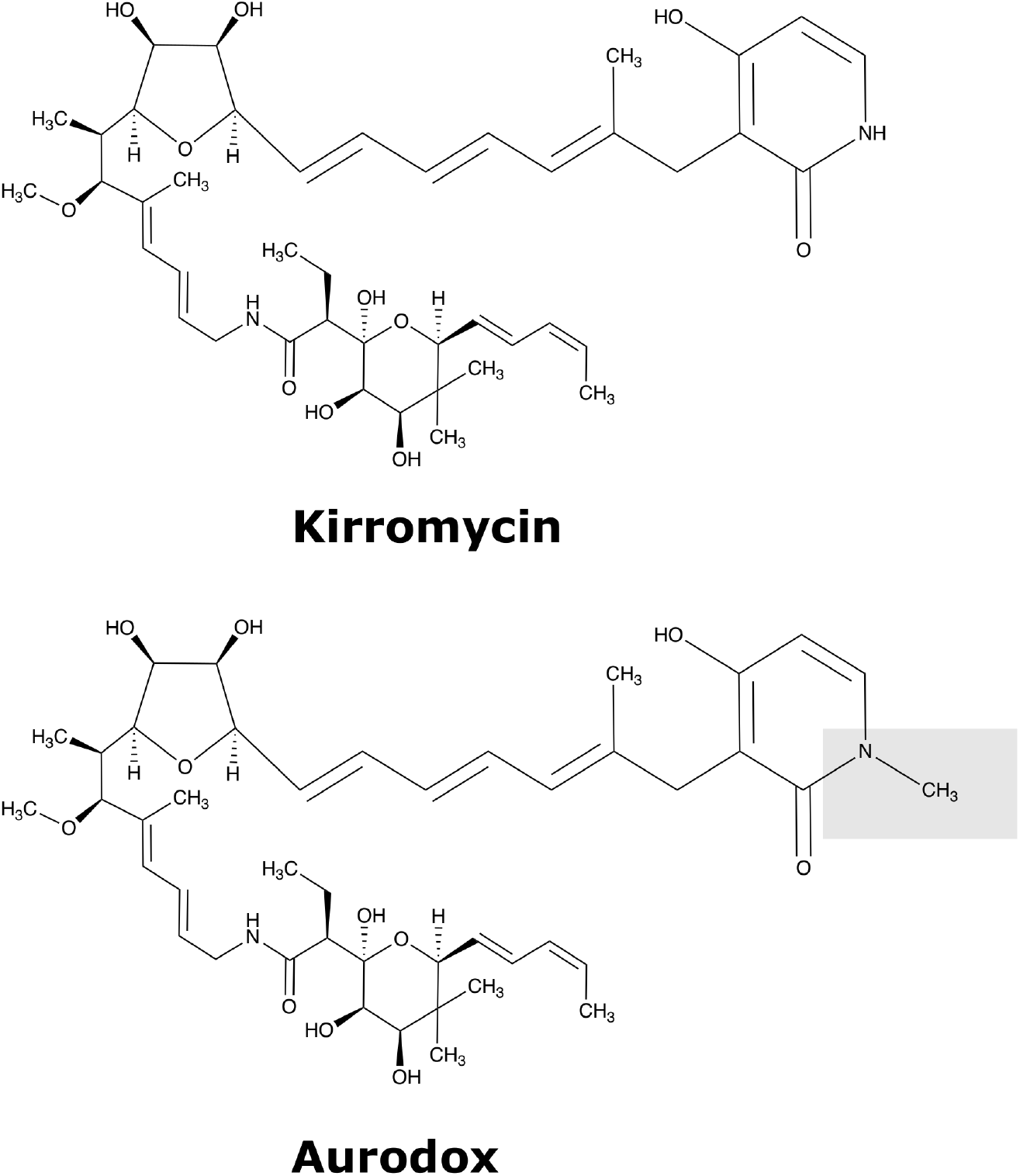
Chemical structures of the elfamycins kirromycin and aurodox. Red box highlights the additional methylation site of aurodox located on the pyridone ring.

In EHEC infections, antibiotic treatment is not recommended due to the prevalence of systemic side effects (11, 12), and the upregulation of the bacterial SOS response in EHEC, resulting in Shiga toxin expression(13–16). The consequences of this are severe for patients with increased Shiga toxin levels being associated with systemic infections and nephrotoxicity (12). Therefore, inhibiting virulence in a specific and targeted manner, that does not induce Shiga toxin production, represents a promising strategy for repurposing aurodox as an anti-virulence treatment (6).

Here, we present the identification and cloning of the aurodox BGC, followed by confirmation of the genes responsible for aurodox biosynthesis through heterologous expression and gene disruption. The aurodox biosynthetic cluster was found to differ in the organisation from that of kirromycin; however, based on homology, we proposed a biosynthetic scheme for aurodox. Additionally, we demonstrate, through heterologous expression of the putative methyltransferase (*aurM\**), that methylation of the pyridone moiety is the last step in aurodox biosynthesis. This improved understanding of aurodox biosynthesis will enable greater exploitation and engineering of aurodox as an anti-virulence therapy and extends our knowledge of an important group of antimicrobial compounds.

## Results

### Whole Genome Sequencing of *S. goldiniensis* reveals a putative aurodox BGC with homology to the kirromycin BGC

To identify the aurodox BGC, the whole genome of *S. goldiniensis* ATCC 21386 was sequenced using a hybrid-approach where Illumina, PacBio and Oxford Nanopore technologies were used to generate a high-quality draft genome (PRJNA602141; (17)). Analysis of the sequence using antiSMASH (18) identified 36 putative BGCs within the *S. goldiniensis* genome **(Supplementary Table S1;** (17). A large region of the *S. goldiniensis* genome was identified (position 4,213,370 - 4,484,508; 271 kb) that was rich in BGCs including an 87 kb region with homology to the kirromycin BGC (7). This 87 kb region consisted of 25 ORFs, with 23 ORFs exhibiting >60% similarity to homologs in kirromycin BGC from *S. collinus* (**Table 1; Supplementary Table S2**). Despite the homologous ORFs within the BGCs, clear differences were apparent between the kirromycin and aurodox gene clusters, such as the inversion of NRPS/PKS genes and rearrangements of genes which encode the decorating enzymes of the polyketide backbone (**Fig. 2**). Two additional genes were identified in the aurodox BGC that lacked homologues in kirromycin cluster (**Table 1**). A gene encoding a SAM-dependant *O*-methyltransferase (*aurM*\*), which we propose catalyses the addition of the methyl group to the pyridone moiety, and a hypothetical protein with no predicted homology to genes of known function (*aurQ*). Given the homology to the kirromycin BGC, it was hypothesised that this putative BGC was responsible for aurodox production in *S. goldiniensis* (**Table 1**).

**Table 1:**
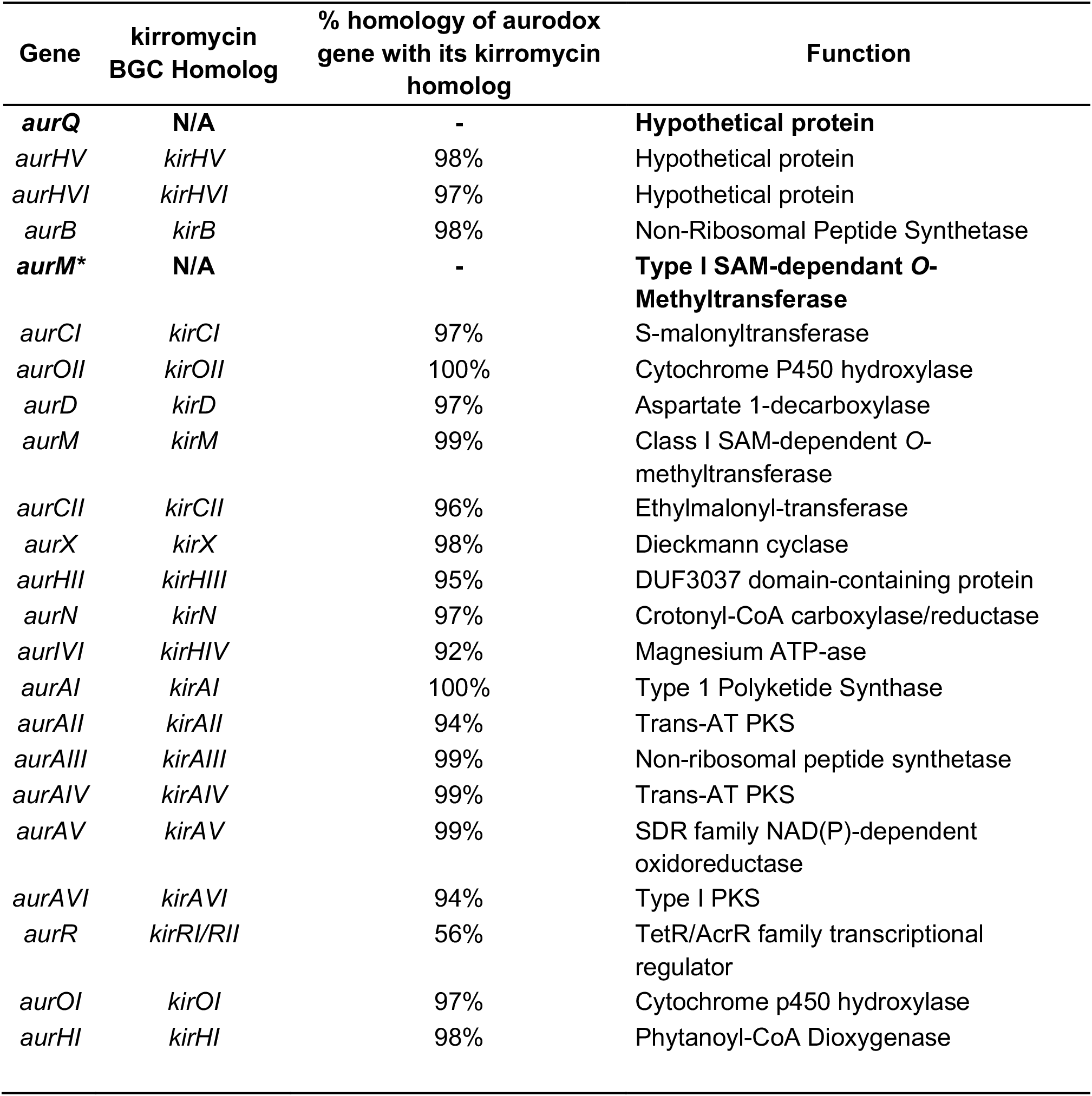
Comparison of aurodox and kirromycin BGC and their functions. Genes found only in the aurodox cluster are in bold. To be consistent with kirromycin biosynthesis we have maintained the nomenclature between this putative aurodox genes and their kirromycin homologs.

**Figure 2:**
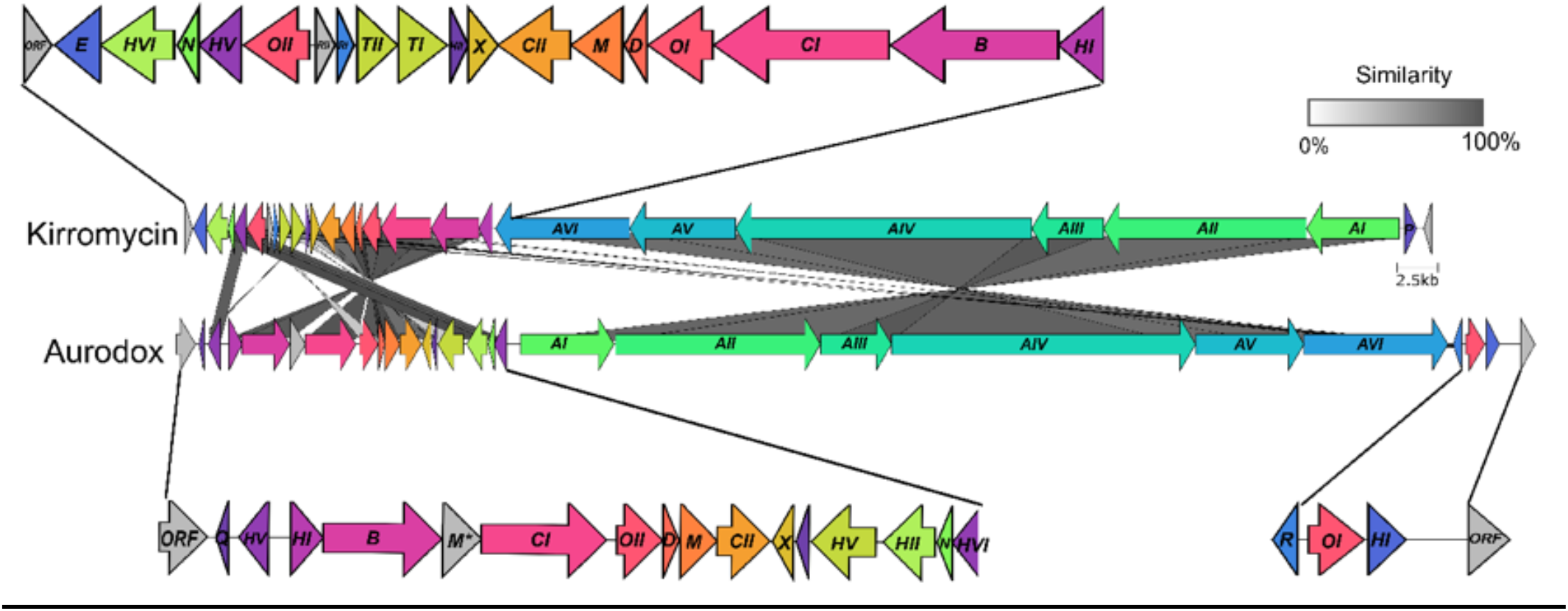
Comparison of aurodox and kirromycin gene clusters. Adjoining lines represent amino acid similarity according to scale. Grey genes represent non-homologous genes, * represents the SAM-dependant *O*-methyltransferase AurM*. Figure generated by clinker.py using (<u>GCA_018728545.1</u>).

### Heterologous expression of the putative aurodox BGC in *Streptomyces coelicolor* M1152 results in aurodox biosynthesis

To determine if the putative aurodox BGC was responsible for aurodox biosynthesis, a Phage Artificial Chromosome (PAC) library was constructed from *S. goldiniensis* genomic DNA, and the resulting pESAC-13A vectors (19) were screened for the presence of the putative aurodox cluster via PCR (Bio S & T, Canada; **Fig. S1;** Oligonucleotide primers can be found in **Supp Table S5**). Two PCR positive PAC clones were identified which contained the entire region of interest, with one clone (pAur1) taken forward for further study (**Fig. S1, S2 & S3**). The complete PAC was sequenced to identify the boundaries of the pAur1, which contained a 129Kbp insert (See **Supp. File. pAur1**; **Fig. S2)** and the complete region proposed to encode the aurodox BGC.

Introduction of pAur1, which integrates at the ɸC31 integration site, to the *S. coelicolor* M1152 ‘superhost’ was achieved via conjugation from the non-methylating ET12567/pR9604 strain to avoid the methyl-specific restriction system (20). Importantly, *S. coelicolor* M1152 encodes three copies of EF-Tu, including one copy of the elfamycin-resistant type EF-Tu, *tuf2* suggesting this strain would be a suitable host for expression of aurodox. Exconjugants containing the putative aurodox BGC (pAur1) and empty vector controls (pESAC-13A) were screened via PCR (**Fig. S3**) and the resulting strains were cultured in liquid media. Culture supernatant extracts were then subjected to LCMS analysis and compared to an authentic aurodox standard (**Fig. 3A)**. An equivalent aurodox peak was also observed in the trace from *S. goldiniensis* extract (**Fig. 3B**). Extracts from the *S. coelicolor* M1152/pESAC-13A, empty vector control lacked the distinct peak of aurodox (**Fig. 3C)**, whereas *S. coelicolor* M1152/pAur1, the strain containing the putative aurodox cluster, exhibited the characteristic peak (**Fig. 3D**). Mass spectrometric analysis revealed peaks at m/z 793, corresponding to the molecular ion of aurodox from cultures of *S. coelicolor* M1152/pAur1 and *S. goldiniensis*. This peak was absent from the empty vector control strain (**Fig. 3C),** indicating that the putative aurodox BGC encodes aurodox biosynthesis in *S. goldiniensis*.

**Figure 3:**
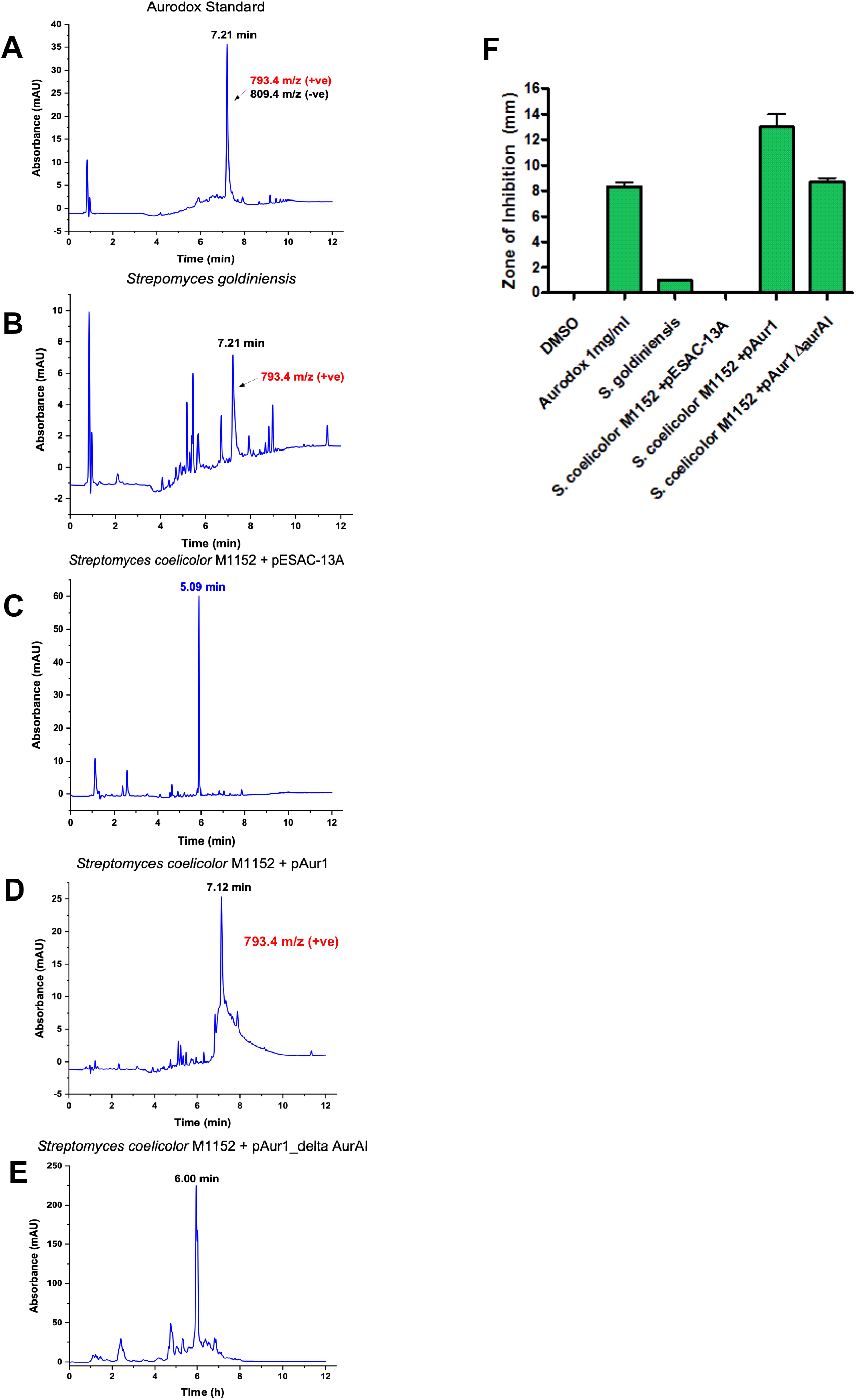
Heterologous expression of aurodox biosynthesis in *Streptomyces coelicolor* M1152. Chromatograms of aurodox standard (Enzo; **A**). Peak at aurodox retention time is indicated. **(B)** Chromatogram from Wild-Type *Streptomyces goldiniensis* indicating aurodox production. **(C)** Chromatogram from the empty vector (pESAC-13A) control strain *S. coelicolor* M1152, indicating the absence of an aurodox-associated peak. **(D)** Chromatogram of extract from growth of *S. coelicolor* M1152/pAur1. Aurodox peak is visible at 7.12 min, and the MS data indicate the presence of aurodox. (**E**) Chromatogram of extracts from *S. coelicolor* M1152/pAur1Δ*aurA*1 showing absence of an aurodox-associated peak. Corresponding MS data can be found in **Fig. S4**. **(E)** Bioactivity of the extracts used in **A-E**, indicating the zones of inhibition associated with the extracts by disc diffusion against *Staphylococcus aureus* ATCC 43300.

To unequivocally confirm the role of the putative aurodox BGC genes encoded on the pAur1 clone, a deletion of the Type IPKS (*aurAI*) was performed using ReDirect (21), resulting in pAur1Δ*aurAI*. We hypothesised that the deletion of the primary PKS would prevent aurodox biosynthesis and confirm the putative BGC was required for aurodox biosynthesis. Analysis of extracts from *S. coelicolor* M1152/pAur1Δ*aurAI* lacked a peak at the aurodox-associated retention time (**Fig. 3E)** confirming the role of these genes in aurodox biosynthesis.

Bioassays of these culture supernatants using *Staphylococcus aureus* (ATCC 43300) as an indicator organism, further support the LCMS data, with *S. coelicolor* M1152/pAur1 inhibiting *S. aureus* growth, whereas the empty vector control (*S. coelicolor* M1152/pESAC-13A) and the deletion strain (*S. coelicolor* M1152/pAur1Δ*aurAI*) display reduced bioactivity (**Fig. 3F)**.

### Proposed biosynthesis of aurodox follows that of kirromycin

The ClusterTools algorithm from antiSMASH was used to annotate the core PKS/NRPS genes of the aurodox BGC including specific module assignments (18, 22). This facilitated the prediction that the catalytic domains of AurI to AurVI largely follow those of KirI to KirVI of the kirromycin gene cluster despite the rearrangements in the overall cluster architecture (**Fig. 2 & 4;** (7, 9). In AurAI and AurAII, acetyl Co-A extension is via Claisen condensation reactions (8), however the PKSs are atypical in arrangement, with two additional dehydratase domains present. Whilst these were not previously identified in the kirromycin pathway(7), reanalysis of the kirromycin pathway using ClusterTools (22) does predict the presence of these domains in KirI to KirVI. Remarkably, no homolog of *kirP,* the kirromycin phosphopanthetheinyl transferase (PPTase) was identified in the aurodox BGC.

It is predicted that *aurAIII* encodes a hybrid NRPS/PKS enzyme consisting of consecutive condensation and adenylation domains which catalyse the condensation of glycine and the incorporation of the amide bond, a process conserved with the kirromycin pathway. The enzymes AurAIV-AVII are predicted to extend the aurodox backbone before AurB (which possesses the conserved DTLQLGVIWK motif (23), catalyses the incorporation β-alanine, presumably synthesised by the putative aspartate-1-decarboxylase, AurD (**Fig. 4 & Table 1)**.

**Figure 4:**
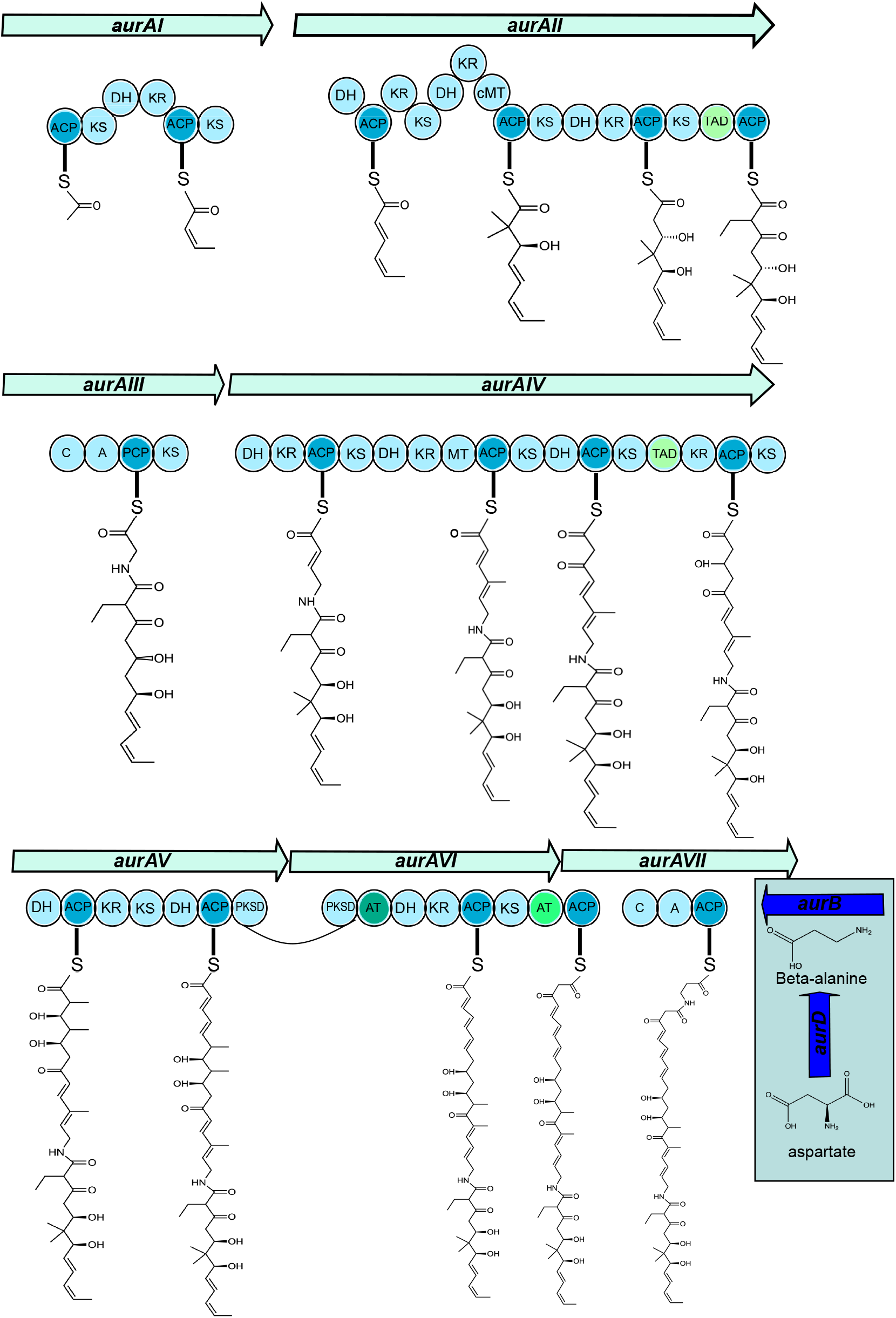
Summary of the enzymatic reactions carried out by AurAI-AurAVII Iin aurodox PKS backbone biosynthesis. ACP: acyl carrier protein; AT: acyltransferase domain; C: condensation domain; DH: dehydratase domain ER: enoyl reductase domain; KR: keto reductase domain; KS: keto synthase domain; MT: methyl transferase domain; PCP: peptidyl carrier protein; TAD: trans-AT docking domain

### A SAM-dependent methyltrasferase, *aurM\** catalyses the conversion of kirromycin to aurodox

An additional SAM-dependant *O*-methyltransferase (AurM*) was identified in the aurodox BGC, which was absent in the kirromycin BGC. Preliminary analysis of the LCMS traces from fermentations of *S. goldiniensis* and *S. coelicolor* M1152/pAur1 indicated that small amounts of kirromycin were present in samples, suggesting that the methylation of kirromycin may be the final step in aurodox biosynthesis. It was hypothesised that AurM* catalyses the conversion of kirromycin to aurodox. To test this, *aurM\** from *S. goldiniensis* was cloned in to an integrating vector (pIJ6902; (24) and introduced in to *Streptomyces collinus* Tü 365, a natural kirromycin producer. Empty vector controls of *S. collinus* Tü 365 containing pIJ6902 showed no species corresponding to aurodox but did show the presence of kirromycin when compared to an authentic standards (**Fig. 5A & 5B)**. LCMS analysis of solvent extracts from *S. collinus* Tü 365 expressing *aurM\** revealed characteristic peaks corresponding to aurodox and kirromycin (**Fig. 5C)**, with negative scan MS showing a m/z ratio of 793, corresponding to aurodox in addition to a m/z ratio 785 which corresponds to kirromycin **(Fig. 5A & Fig S5)**. This indicates that AurM* is responsible for the methylation of kirromycin as a precursor to aurodox formation.

**Figure 5:**
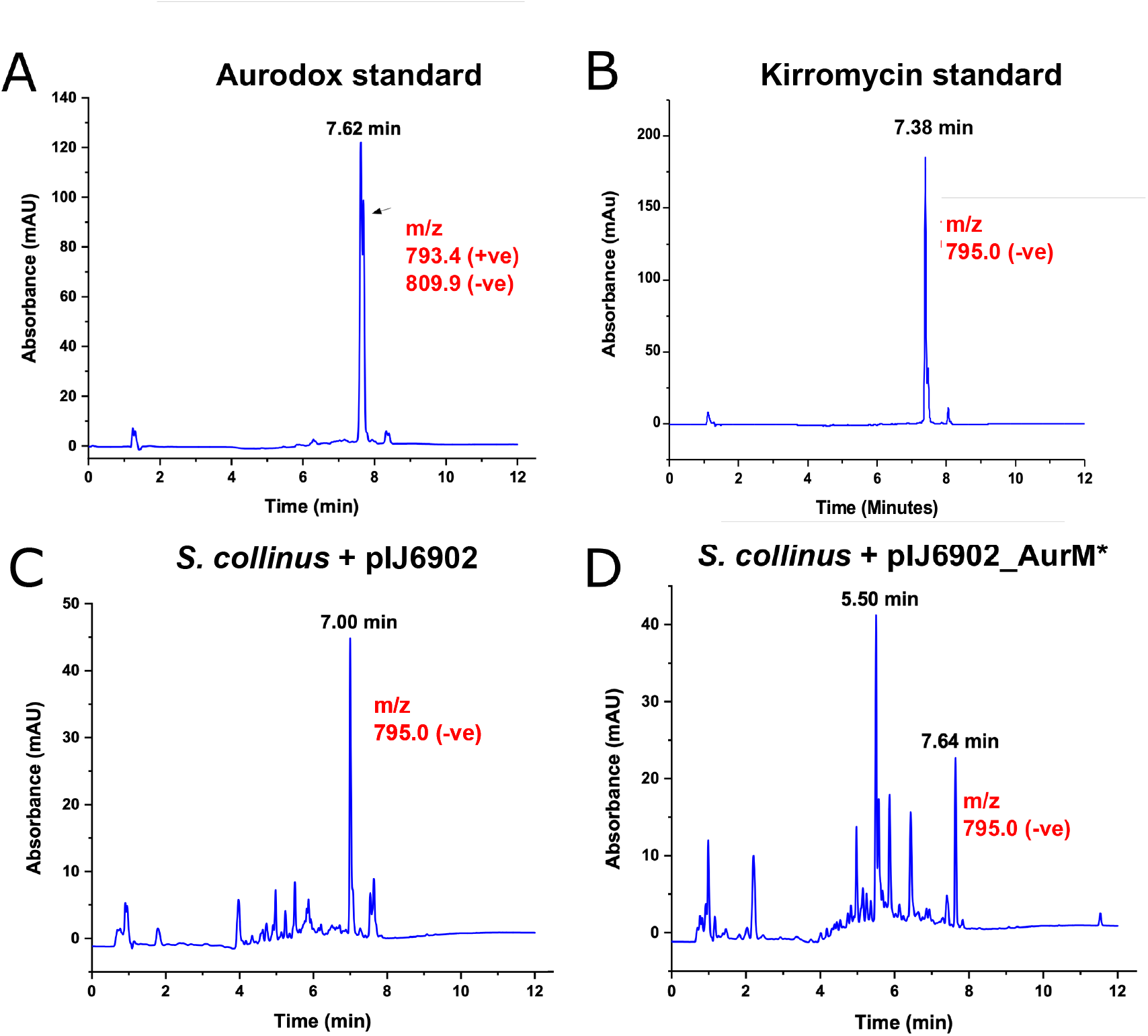
Methylation of kirromycin is the last step in aurodox biosynthesis and is catalysed by AurM*. Chromatograms of aurodox **(A)** and kirromycin **(B)** standards and fermentation extracts from a control strain *S. collinus* Tü 365 containing pIJ6902 (empty vector control; **C**) showing presence of kirromycin and absence of aurodox-associated mass and **(D)** chromatogram from extracts from *S. collinus* Tü 365 containing pIJ6902_AurM* showing the presence of aurodox with a retention time of 7.64 min. Corresponding MS data can be found in **Fig. S5**.

## Discussion

Understanding the mode of action, resistance mechanisms and the biosynthesis of useful natural products is key to their development for clinical application. The linear polyketide, aurodox, whilst being known for almost 50 years, was recently found to exhibit novel anti-virulence activity via a previously unknown target (6), yet nothing was known about its biosynthesis. Whilst aurodox is structurally similar to kirromycin, it is now well known that structurally identical or highly similar natural products can be biosynthesised via diverse chemical routes (1, 25–28), suggesting that there is still much to be learned from studying biosynthesis of structurally similar natural products in terms of novel activity and evolution of natural products.

Despite similarities in structure, anti-bacterial mode of action and core BGC machinery between aurodox and kirromycin, key differences in biosynthesis were identified. The aurodox cluster is ~80% similar to the kirromycin BGC, sharing 23 of the 25 genes, with seven genes within both clusters encoding hypothetical proteins with no assigned function. There is no apparent PPTase (*kirP* homolog) encoded in the aurodox cluster, which would normally post-translationally modify the acyl carrier protein (ACP) to facilitate extension of the PK/NRP chains during assembly, suggesting that this function may be fulfilled by a promiscuous PPTase encoded elsewhere in the genome (29). Remarkably, no thioesterase domain was identified in the PKS-NRPS megasynthases. There is a putative Dieckmann-like cyclase encoded within the cluster (*aurB*), which may be responsible for the cleavage and cyclisation of the aurodox molecule, a mechanism that recently been proposed in a few other pyridone natural products (30). The anti-virulence activity of aurodox requires methylation of the pyridone ring, which is catalysed by *aurM\**. The ‘magic methyl’ effect is well known in drug discovery for enhancing the activity and pharmacological properties of drugs (31, 32), and the kirromycin to aurodox transformation indicates that this has been exploited in nature to alter the activity of natural products. Moreover, this indicates that derivatisation of the pyridone ring of kirromycin may be a useful strategy for diversifying elfamycin activities. Regarding the origin of *aurM*,* two additional *O*-methyl transferases are encoded within the *S. goldiniensis* genome, each with ~35% amino acid similarly to AurM*, which may suggest that *aurM\** was acquired by horizontal gene transfer (HGT) rather than duplication of an existing *O*-methyltransferase from the genome of *S. goldiniensis,* and offers insight in to the evolution of novel activities in natural products and the potential of this to be driven through HGT of single ORFs.

Overall, this greater understanding of biosynthesis of aurodox and the steps that contribute to unique modes of action will enable us to explore the potential of aurodox as a therapeutic agent for EHEC treatment in the food chain and the clinic.

## Materials and Methods

### Growth and Maintenance of Bacterial Strains

The bacterial strains used in this study are detailed in **Table S3**. Genetic constructs used and their antibiotic selection are described in **Table S4**. Routine growth and maintenance procedures were carried out according to Kieser *et al,*(33). A list of oligonucleotides used in this study can be found in **Table S5**.

### Whole Genome Sequencing *Streptomyces goldiniensis*

Genomic DNA was extracted from *S. goldiniensis* using the Streptomyces DNA isolation protocol described by Kieser *et al,* (33). Nanopore reads were obtained using a genomic DNA library prepared in accordance with the Nanopore™ 1D ligation protocol, using MinION SPOT ON MK1 R9© flow cells and the raw data was converted to sequence data via MinKNOW base calling software. Illumina data was obtained from Microbes NG (Birmingham, UK) using the HiSeq 2500 sequencing platform. Reads were adapter trimmed using Trimmomatic 0.30 (34) with a sliding window quality cut off of Q15. PacBio sequencing was provided by Nu-omics at (University of Northumbria, UK) using the PacBio Sequel instrument and contigs were assembled in HGAP4.

*Streptomyces goldiniensis* genome was assembled using SPAdes (34) using data provided by all three platforms. AutoMLST (35) was used to determine the closest neighbour *S. bottropensis* ATCC 25435 (Taxonomy ID: 1054862) which was used for scaffold-based assembly via MeDuSa(36), with quality analysis carried out using QUAST (37). Annotation of the *Streptomyces goldiniensis* genome was created using Prokka (38) and can be accessed at the Genbank Bioproject PRJNA602141. Biosynthetic gene cluster identification was carried out using antiSMASH bacterial version 5.0.0 with modular enzymatic domains analysis carried out using the PKS/NRPS domain annotation function in antiSMASH (18).

### Aurodox production, purification and detection

*Streptomyces* spore stocks (1 × 10^8^ spores) were pre-germinated in 10 ml of GYM medium (32) overnight at 30 °C with shaking at 250 rpm. Cells were then washed three times to remove antibiotics (if used) and resuspended in 1ml of GYM, which was used to inoculate a 50 ml seed culture of GYM which were incubated at 30 °C with shaking at 250 rpm. After seven days, biomass was removed by centrifugation (4000 × *g*, 10 minutes) and culture supernatant was filter sterilised through a 0.2 μM Millipore™ filter. Supernatants were mixed with equal volume of chloroform and a separation funnel was used to remove the lower, solvent phase. Solvent extracts were dried under nitrogen and extracts were solubilised in ethanol for LC-MS analysis. Authentic aurodox standard (1 mg/ml) was purchased from Enzo (New York, USA). LC-MS was carried out on an Aglient 1100 HPLC instrument in conjunction with a Waters Micromass ZQ 2000/4000 mass detector. Electrospray ionization (ESI) was used in all cases. The RP-HPLC analysis was conducted on a Zorbax 45mm × 150mm C18 column at 40°C. Ammonium acetate buffers were used as follows: Buffer A (5 mM Ammonium acetate in Water) and Buffer B (5mM Ammonium acetate in acetonitrile). Positive and negative electrospray methods were applied with of 100 to 1000 AMU positive, 120-1000 AMU negative with a scanning time of 0.5 seconds. The UV detection was carried out at 254 nm.

### Construction of aurodox expression strains and deletion mutant

Aurodox encoding vector pAur1 (Bio S & T, Canada) and empty vector pESAC-13A (39) were transferred to the non-methylating *Escherichia coli* strain ET12567 via tri-parental mating. Briefly, fresh overnight cultures of *E. coli* DH10β containing pAur1 or the parental vector pESAC-13A (apramycin resistant), *E. coli* Top10 containing the driver plasmid pR9604 (Beta-lactam resistant) and ET12567 (Chloramphenicol resistant) were grown to an OD_600_ of 0.6. Cells were washed three times in fresh LB by centrifugation at 4000 × *g* and resuspended in 1 ml of LB. A 20 μl aliquot of each strain was plated in the centre of a LB plate and incubated at 37 °C overnight. The resulting growth was re-streaked on to the required antibiotic selection and the presence of the conjugating vector in the *E. coli* ET12567/pR9604 strain was confirmed via colony PCR. Mutation in pAur1 were carried out according to the ReDirect PCR targeting system in *Streptomyces* (21), using the hygromycin resistance cassette, pIJ10700 as the disruption cassette (http://streptomyces.org.uk/redirect/RecombineeringFEMSMP-2006-5.pdf). Disrupted PACs were introduced in to *E. coli* ET12567/pR9604 via tri-parental mating and subsequently moved into *Streptomyces coelicolor* M1152 via conjugation as described above.

### Cloning of *aurM\** from *Streptomyces goldiniensis*

The putative *O*-methyltransferase *aurM\** was amplified from *S. goldiniensis* genomic DNA using the oligonucleotide primers in **Table S5** and cloned in to pIJ6902 using the NEB Gibson Assembly cloning kit (24). Conjugation of pIJ6902 based vectors was according to Kieser et al., (33).

### Bioassays

Bioassays were conducted using disc diffusion assays with *Streptomyces* fermentation extracts on soft nutrient overlays containing *Staphylococcus aureus* ATCC43300 as the indicator organism.

## Acknowledgements

We would like to thank Professor Wolfgang Wohlleben (University of Tübingen) for the gift of the *Streptomyces collinus* Tü 365 strain, Dr Margherita Sosio (Naicons) for the gift of pESAC-13A and Professor Mervyn Bibb (John Innes Centre) for the gift of the *S. coelicolor* M1152 strain. We would also like to thank Professor Iain S. Hunter (University of Strathclyde) and Professor Matt Hutchings (John Innes Centre) for helpful discussions.

